# Arnav: Site specific error models to identify variants in RNA

**DOI:** 10.1101/397539

**Authors:** Avinash Ramu, Donald F. Conrad

## Abstract

**Summary:** We present Arnav (Analysis of RNA variants) a lightweight and easy-to-use statistical method for detecting mutations from RNA sequencing data. Site-specific error models allow Arnav to call variants with high specificity when the true variant allele fraction is unknown. We show the utility of Arnav by identifying variants using RNA sequencing data from the GTEx project.

**Availability and Implementation:** Arnav is implemented in C++ and is distributed under the GPL license at https://github.com/gatoravi/arnav.

---

Mutations have the power to specify the course of a genome. Detecting new mutations specific to a sample of interest can help further our understanding of the mutational process and identify the molecular mechanisms that are active or disrupted in a sample. Identifying mutations from bulk tissue samples is technically challenging when the mutation is present in only a small fraction of cells. In this paper we describe a lightweight and easy-to-use statistical method, Arnav (Analysis of RNA Variants), that can be used to detect sequence variants that are present in varying unknown proportions within a sample. The statistical model underlying Arnav is powerful in the setting where a large number of samples have been sequenced using similar library preparation methods. We demonstrate, by analysis of RNA sequencing data, that the Arnav model allows us to detect low frequency variants with high specificity.

## Methods

Arnav constructs a site-specific null expectation for the variant allele fraction (VAF) for each position of interest, *k*, by tabulating base counts from the BAM files of a control cohort. We expect the resulting estimate of the alternate allele fraction at each site, *e*_*k*_, to reflect site-specific errors related to library preparation, sequencing and mapping, but, depending on the source of input material (DNA or RNA), will also incorporate biological sources of variation such as germline variation, allele-specific expression, and RNA editing. In order to identify sequence variants in the test sample, Arnav calculates the probability of sampling the observed read count data at position *k* assuming a binomial model parameterized by the estimated error rate *e*_*k*_. The key advantages of the Arnav statistical model are a) the ability to call variants from diverse sequencing assays such as RNA, ATAC or CHIP sequencing data, b) the simplicity and resulting speed of the method makes it feasible to perform genome-wide or transcriptome-wide calculations on 10,000s of samples, c) the use of site-specific error models can, in favorable sequence contexts, allow sensitivity to mosaic variants that are rarer than the average sequencing error rate.

## Results

To assess the sensitivity of Arnav on calling mutations from RNA sequencing data we constructed site-specific null models using 9597 BAM files from release version 6 of the GTEx project [1]. The distribution of the estimated null error rate, *e*_*k,*_, across sites in chromosome 22 is shown in Fig. 1a and it varies by orders of magnitude across all sites (min:5.6e-9, median:4.5 e-4, max: 1). This heterogeneity in *e*_*k*_confirms the utility of site-specific error models for mutation calling in this setting. We observe an 80-fold increase in the median alternate allele fraction at known RNA editing sites(min = 6.22e-5, median: 0.036, max: 1) from RADAR[6]. This enrichment highlights the ability of the error model to capture real biological signal.

Next, we used the estimated null models to call mutations using RNAseq data from blood samples of five GTEx individuals (Figure 1b). A fraction of the variants called by Arnav overlapped the germline DNA variants identified in the same individuals by the GTEx consortium; these are plotted in a separate panel. The sites that remain after filtering against germline variants are expected to include putative post-zygotic DNA mutations, rare post-transcriptional modifications and technical artefacts.

We used two alternative variant calling approaches to benchmark variant calling using Arnav. We first ran the GATK best practices for variant calling on RNA-seq pipeline[7][8] on the same input data. We also implemented a naive read-count based approach to compare with results from model based approaches. In the naive approach we call as a variant any site with greater than five reads of support for the variant allele and VAF greater than 10%. Across five samples, Arnav identified a mean of 6487 variants per sample (median: 6512), GATK identified a mean number of 83976 variants per sample (median: 81352) and the naive variant calling approach identified a mean number of 26972 variants per sample (median: 22799).

We next assessed the specificity of the three variant calling approaches. We attempted to in-silico validate variant calls with greater than five reads of support and VAF greater than 10%. To do this we used exome sequencing data derived from the blood of the same five individuals. We marked as validated any site in the exome data with two or more reads supporting the variant allele identified by the original variant caller. The validation rate was calculated as the number of validated sites divided by the total number of putative variant sites with read support in the exome data. Note that this validation rate does not penalize variant calls with an absence of read coverage in the exome data. Arnav’s site-specific error modeling resulted in a mean validation rate of 86% (mean number of validated calls: 1243, median: 1258) compared to a validation rate of 56% using GATK(mean number of validated calls: 3783, median: 3873) and a validation rate of 20% using the naive variant caller (mean number of validated calls: 1745, median: 1762.) The validation rates for the individual samples are shown in Figure 1c. While a greater number of calls are validated using GATK and the naive variant caller, the fewer variant calls and the resulting increase in the positive predictive value using Arnav makes it more practical for variant calling. The VAF in the RNA sequencing data and the exome data at validated sites called by Arnav are plotted in Figure 1d. Many of the sites identified by Arnav are putative germline heterozygous or homozygous variants but we also see validation support in the exome data for variants identified by Arnav across other regions of the allele frequency spectrum.

In addition to the statistical model, we have identified downstream filtering steps that aid in removing mutation artefacts not captured by the error model. These steps are included in the software repository as a BASH script that users may modify depending on their study design. For example, users interested in somatic mutations may want to filter out known RNA editing sites. We have implemented four general filters:

1. Require a minimum number of reads supporting the variant allele (we use a minimum of five reads).
2. Require reads supporting the variant allele on both strands.
3. If available, remove germline polymorphisms from the same sample.
4. Check for bias in the position of the variant allele within the reads. This is done using the VDB metric implemented in samtools[4].

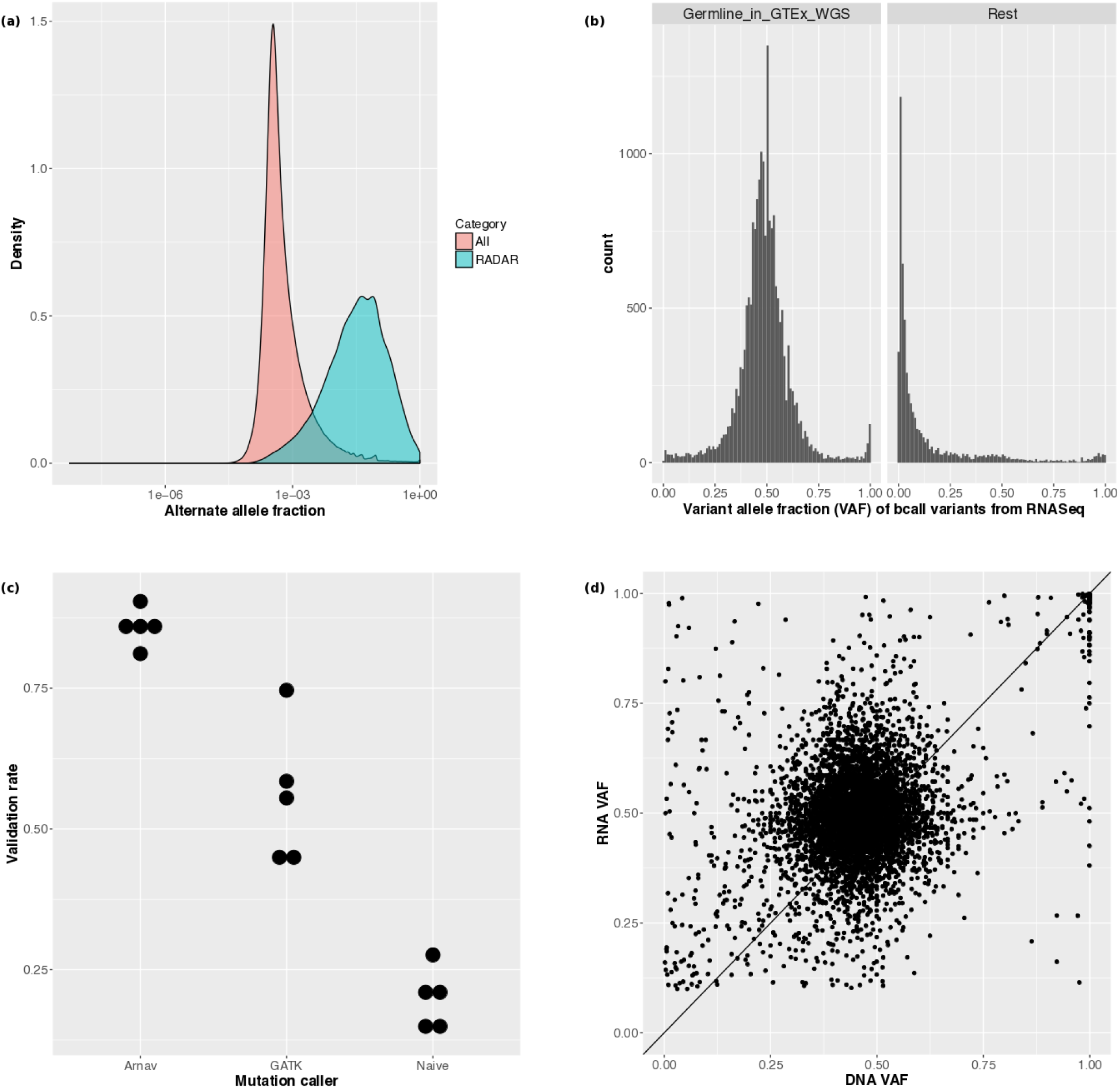

ARNAV can be used to reliably identify mutations from RNA sequencing data. (**a**) Distribution of non-reference allele fraction, plotted on a log scale, across chr22 sites (n = 3,524,378) in all samples. Sites with at-least one read supporting the non-reference allele are plotted. This distribution summarizes the site-specific null error models across all sites. Known RNA-editing sites in chr22 (n = 19,579) from RADAR[6] are plotted separately and show a higher non-reference allele fraction reflecting likely RNA editing signal. (**b**) Distribution of VAF at sites called by Arnav in five blood RNA sequencing samples. Sites overlapping germline variants identified by the GTEx consortium using Whole Genome Sequencing data are plotted separately from the rest of the variant calls. (c) Validation rate of variants identified in blood derived RNA from five individuals in the GTEx cohort. Variants were identified using Arnav, GATK, and using a naive read-count based variant caller. Variant calls with at least two reads of support for the variant allele in the exome data were marked as validated. (d) VAF of the validated variant calls from Arnav in the blood RNA sequencing data plotted against the VAF in the exome data. Only sites with more than ten reads in both datasets are shown.

## Implementation

Arnav is implemented in C++ and is distributed under the GPL license at https://github.com/gatoravi/arnav. Arnav uses open source libraries from the R project[1] and cereal[2]. Read-counts for each site in the genome are extracted using mpileup2readcounts[3] and requires samtools[4] to be installed.

## Acknowledgements

We thank David E. Larson for feedback on the manuscript and Nicole Rockweiler for feedback on Arnav. This work was supported by National Institutes of Health Grants R01HD078641 and R01MH101810 to D.F.C. Washington University holds an Institutional Program unifying Population and Laboratory Based Sciences Award from the Burroughs Wellcome Fund. AR is supported by this program.

